# A Semantic + Neuronal Approach to Predict Pathogenic Variants in DNA Sequences

**DOI:** 10.64898/2026.08.16.745093

**Authors:** Jesus Antonio Motta, Carolina Fernandez, Maria del Mar Motta

## Abstract

In this work, we present a machine learning model for identifying pathogenic DNA variants. The model was learned from the analysis of normal and pathogenic sequences extracted from the ClinVar database (supported by NCBI). This analysis was based on a conceptual semantic model of DNA sequences converted to peptide sequences (amino acid sequences) governed by a well-defined grammar, which allowed us to apply NLP techniques, specifically Part of Speech tagging (POS tagging). Our predictive model was built by combining two techniques: CRF (from the Markov model family), which performs the sequencing, and BiLSTM (a deep learning model) which captures the past and future content of the sequences. The training space was created with the sequences of 102 genes associated with approximately 27,000 pathogenic variants. The model was evaluated using the metrics precision, P-R and ROC curves, AUC, and confusion matrices. Its performance was also compared against five known methods for predicting pathogenic variants. The results show exceptional performance that exceeds expectations and places this new method at the state of the art for predicting pathogenic DNA sequences.

**Rationale:** Biological sequences exhibit a **hierarchical organization** that parallels the structure of natural language. At the residue level, amino acids function as **functional tokens**: catalytic residues behave like **verbs** that drive biochemical actions, hydrophobic residues form **noun-like structural cores**, and regulatory residues act as **modifiers** that tune activity. Short sequence motifs correspond to **phrase-like units**, domains operate as **clause-level structures**, and full proteins form coherent **sentences** within the broader **paragraphs** of cellular pathways. This linguistic analogy provides a natural justification for applying **semantic and syntactic modeling frameworks** to protein sequences.

By treating sequence elements as tokens embedded within a **grammar-like system**, we can analyze variant effects, domain interactions, and regulatory motifs using tools originally developed for **natural language understanding**. Conceptualizing proteins as linguistic constructs enables the use of **semantic embeddings**, **contextual encoders**, and **hierarchical attention mechanisms** to capture dependencies among residues, motifs, and domains that are not apparent from primary sequence alone. This perspective also offers a principled framework for interpreting model behavior: attention to serine or tyrosine residues corresponds to **verb detection**, clustering of hydrophobic residues reflects **noun-like structural cores**, and interactions among domains map to **clause-level syntax**.

In natural language, meaning arises not only from individual words but from their **grammatical relationships**, **contextual dependencies**, and **hierarchical composition**. Proteins exhibit analogous properties. Catalytic residues act as verbs that initiate biochemical reactions; hydrophobic cores provide the structural nouns; regulatory residues function as modifiers; and flexible linkers serve as conjunctions that connect functional units. Domains behave as clauses whose interactions determine the overall **syntactic behavior** of the protein. Mutations disrupt this grammar in predictable ways—ranging from **minor spelling changes** (missense variants) to **truncated sentences** (nonsense mutations) and **syntactic collapse** (frameshifts). This linguistic framework therefore supports the development of **interpretable, biologically grounded machine-learning models** capable of capturing the multi-level dependencies that govern protein function and variant impact.

## 1. Introduction

Accurate interpretation of genetic variants remains one of the central challenges in human genomics. Although large-scale sequencing has identified millions of rare variants across human populations, only a small fraction has well-established clinical significance. The majority remain classified as variants of uncertain significance (VUS), limiting their utility in diagnosis, risk prediction, and precision medicine. Computational prediction methods have therefore become essential tools for prioritizing potentially pathogenic variants and guiding downstream experimental or clinical evaluation.

Despite these advances, deep learning models for variant pathogenicity remain underdeveloped relative to their success in regulatory genomics. Existing pathogenicity predictors often rely on limited sequence context, do not incorporate long-range interactions, or fail to model structured dependencies across genomic regions. Moreover, many models are optimized for specific variant types—such as missense substitutions—leaving indels and noncoding variants comparatively underserved. These limitations highlight the need for new architectures that integrate local sequence motifs, long-range contextual information, and structured dependencies to improve pathogenicity prediction across diverse variant classes.

Early approaches to variant pathogenicity prediction relied on evolutionary conservation, protein structural features, and manually engineered annotations. Representative methods include **SIFT** [1], which evaluates amino acid substitutions based on evolutionary constraint, and **PolyPhen-2** [2], which integrates sequence, structural, and comparative genomics features to estimate functional impact. Ensemble-based predictors such as **CADD** [3] and **REVEL** [4] further advanced the field by combining diverse annotations into unified scores that correlate with pathogenicity. These tools remain widely used in clinical genomics pipelines, yet they are limited by their dependence on predefined features and their inability to fully capture the complex sequence context surrounding a variant.

In this work, we introduce a deep learning framework that combines convolutional, recurrent, and structured related sequential components to model variant effects directly from genomic sequence. We evaluate our approach against widely used predictors, including **CADD**[3]**, SpliceAI**[5]**, gnomAD-AF**[6]**, Strvctvre**[7] and **Expansion Hunter**[8] as well as state-of-the-art deep learning models, demonstrating improved performance across multiple variant categories. Our results illustrate the potential of hybrid architectures to advance variant interpretation and contribute to more accurate genomic medicine.

## 2. Related Work

Deep learning has transformed computational genomics by enabling models to learn hierarchical representations directly from raw biological sequences. Convolutional neural networks (CNNs) [9], [10] have demonstrated strong performance in predicting chromatin accessibility, transcription factor binding, and regulatory variant effects, as exemplified by **DeepSEA**[11], **Basset**[12], and **Basenji**[13] (Kelley et al., 2018). Hybrid architectures that combine CNNs with recurrent neural networks (RNNs), such as **DanQ**[14], have shown that long-range dependencies in DNA can be captured more effectively by integrating convolutional motif detectors with bidirectional LSTMs. Structured prediction frameworks, including **Conditional Random Fields (CRFs)** [15], have further been applied to genomic segmentation tasks, demonstrating the value of modeling dependencies between adjacent sequence positions. Table 1 presents a comparison of variant pathogenicity prediction methods where can be observed the type or class of model, paper source, metric type, typical performance, main strengths and weaknesses and model type among five: statistical/rule based, classical machine learning (ML), deep learning, evolutionary/generative models and non-ML/algorithmic.

**Table 1.**
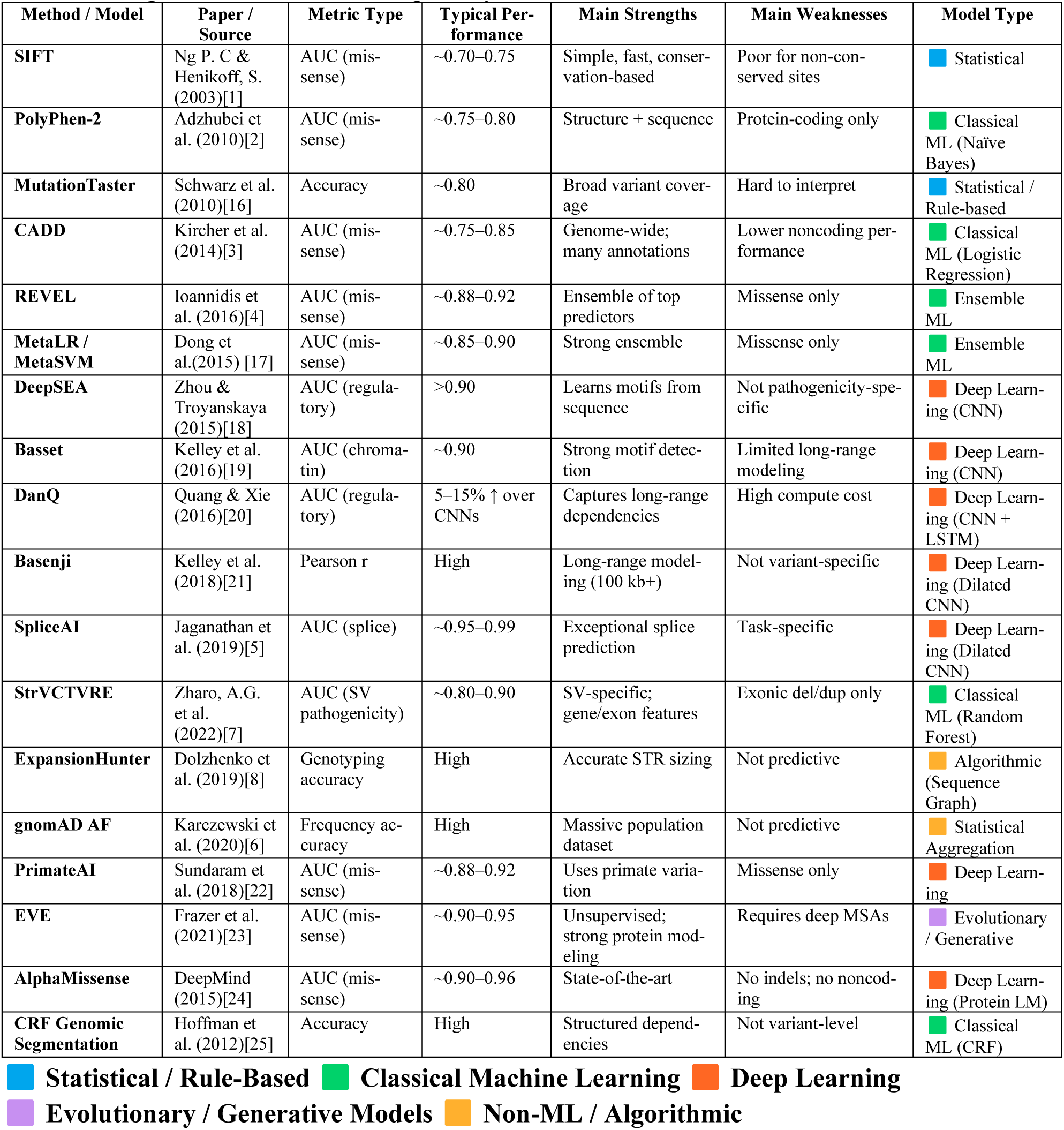
Comparison of Variant Pathogenicity Prediction Methods.

| Method / Model | Paper / Source | Metric Type | Typical Performance | Main Strengths | Main Weaknesses | Model Type |
| --- | --- | --- | --- | --- | --- | --- |
| **SIFT** | Ng P. C & Henikoff, S. (2003)[1] | AUC (mis-sense) | ~0.70–0.75 | Simple, fast, conservation-based | Poor for non-conserved sites | Statistical |
| **PolyPhen-2** | Adzhubei et al. (2010)[2] | AUC (mis-sense) | ~0.75–0.80 | Structure + sequence | Protein-coding only | Classical ML (Naïve Bayes) |
| **MutationTaster** | Schwarz et al. (2010)[16] | Accuracy | ~0.80 | Broad variant coverage | Hard to interpret | Statistical / Rule-based |
| **CADD** | Kircher et al. (2014)[3] | AUC (mis-sense) | ~0.75–0.85 | Genome-wide; many annotations | Lower noncoding performance | Classical ML (Logistic Regression) |
| **REVEL** | Ioannidis et al. (2016)[4] | AUC (mis-sense) | ~0.88–0.92 | Ensemble of top predictors | Missense only | Ensemble ML |
| **MetaLR / MetaSVM** | Dong et al. (2015) [17] | AUC (mis-sense) | ~0.85–0.90 | Strong ensemble | Missense only | Ensemble ML |
| **DeepSEA** | Zhou & Troyanskaya (2015)[18] | AUC (regulatory) | >0.90 | Learns motifs from sequence | Not pathogenicity-specific | Deep Learning (CNN) |
| **Basset** | Kelley et al. (2016)[19] | AUC (chromatin) | ~0.90 | Strong motif detection | Limited long-range modeling | Deep Learning (CNN) |
| **DanQ** | Quang & Xie (2016)[20] | AUC (regulatory) | 5–15% ↑ over CNNs | Captures long-range dependencies | High compute cost | Deep Learning (CNN + LSTM) |
| **Basenji** | Kelley et al. (2018)[21] | Pearson r | High | Long-range modeling (100 kb+) | Not variant-specific | Deep Learning (Dilated CNN) |
| **SpliceAI** | Jaganathan et al. (2019)[5] | AUC (splice) | ~0.95–0.99 | Exceptional splice prediction | Task-specific | Deep Learning (Dilated CNN) |
| **StrVCTVRE** | Zharo, A.G. et al. (2022)[7] | AUC (SV pathogenicity) | ~0.80–0.90 | SV-specific; gene/exon features | Exonic del/dup only | Classical ML (Random Forest) |
| **ExpansionHunter** | Dolzhenko et al. (2019)[8] | Genotyping accuracy | High | Accurate STR sizing | Not predictive | Algorithmic (Sequence Graph) |
| **gnomAD AF** | Karczewski et al. (2020)[6] | Frequency accuracy | High | Massive population dataset | Not predictive | Statistical Aggregation |
| **PrimateAI** | Sundaram et al. (2018)[22] | AUC (mis-sense) | ~0.88–0.92 | Uses primate variation | Missense only | Deep Learning |
| **EVE** | Frazer et al. (2021)[23] | AUC (mis-sense) | ~0.90–0.95 | Unsupervised; strong protein modeling | Requires deep MSAs | Evolutionary / Generative |
| **AlphaMissense** | DeepMind (2015)[24] | AUC (mis-sense) | ~0.90–0.96 | State-of-the-art | No indels; no noncoding | Deep Learning (Protein LM) |
| **CRF Genomic Segmentation** | Hoffman et al. (2012)[25] | Accuracy | High | Structured dependencies | Not variant-level | Classical ML (CRF) |

**Statistical / Rule-Based** **Classical Machine Learning** **Deep Learning** **Evolutionary / Generative Models** **Non-ML / Algorithmic**

## 3. Theoretical bases of used methods

### 3.1 Conceptual Metaphor

Considering the fascinating analogy between a genetic sequence and a complex semantic unit (for example, a sentence), where we can identify nucleotides (A, T, G, C) as symbols (letters), codons (triplets of nucleotides) as words, and genes formed with sequences of words (sentences), which have biological meaning by producing a protein. Promoters and enhancers can be likened to punctuation, guiding when and how the sentence is read. Furthermore, a mutation can be interpreted as a typo or grammatical shift that alters the meaning or disrupts the function. Thus, we can consider a genetic sequence, using a conceptual metaphor, as a complex semantic unit with special characteristics:

– Context. Just as a word can mean different things in different sentences, a codon or gene can behave differently depending on its genomic context.
– Redundancy and Ambiguity. Multiple codons can code for the same amino acid (synonyms), and a mutation can be silent or disruptive, like homonyms or spelling errors.
– Grammar. The grammar of gene expression involves enhancers, silencers, and epigenetic markers for tone, emphasis, or sentence structure.

The preceding concepts and analogies allow us to characterize them as a whole in a conceptual metaphor rather than a literal mapping, in the sense that biology does not assign POST to amino acids; however, their functional roles, motifs, and domains behave in a way that resembles a linguistic structure, as we have mentioned previously.

To illustrate these ideas in relation to the parts of a sentence, we would have:

– Nouns. These are the structural amino acids that provide substance or identity to a protein, such as:

a. Leucine, Isoleucine, Valine. These are the “core-builders” that form the protein’s body.
b. Glycine. Small and flexible, it is a “pivot point”
c. Proline. A rigid “stop sign” that breaks helices

These amino acids behave like nouns: they define what the protein is made of

– Verbs. These are the catalytic or regulatory amino acids. These amino acids do things, they perform actions. For example:

a. Serine, Threonine, Tyrosine, which control the phosphorylation sites, switching the activity on and off
b. Histidine, Cysteine, Aspartate. This is a catalytic triad in the enzyme TEV (It is a specific cysteine protease that has several applications, including the removal of fusion tags in recombinant proteins). A catalytic triad is a set of 3 coordinated amino acids that can be found in the active site of some enzymes
c. Lysine. It acts in ubiquitination and acetylation, which are the signals for degradation/activation.

These amino acids behave like verbs, combining the state of the protein or cell.

– Adjectives. Some amino acids modify the interaction of a protein with its environment. Examples:

a. Valine, Leucine. They increase hydrophobicity. They change the membrane association.
b. Aspartate, Glutamate. They add a negative charge. They also alter the affinity for binding.
c. Arginine, Lysine. They facilitate the positive charge for DNA binding.

These behave like adjectives: they modify how the protein behaves.

– Adverbs. They are modifiers of context dependence. Some residues do not act directly, but influence how strongly or under what conditions other residues (amino acids in enzymes that play crucial roles facilitating biochemical reactions) act. For example:

a. Proline. Near a phosphorylation site, it enhances kinase recognition.
b. Glycine. Near catalytic residues, it increases flexibility and modulates the reaction rate.

The above amino acids behave like adverbials: they modify when and how an action occurs.

– Conjunctions. They are linkers: Flexible or neutral residues connect functional units. Examples:

a. Gly-Ser repeats in flexible linkers
b. Poly-alanine stretches in transcription factors

They behave like conjunctions connecting clauses (domains).

– Punctuation. Stop codons and structural breaks. Examples:

a. Stop codons: periods
b. Signal peptides: commas (pause before secretion)
c. Proline. Helix Breaker. As a semicolon

Table 2 shows a mapping of all 20 amino acids to linguistic roles (Functional Tendencies), table 3 presents a formal annotation tagset and figure 1 shows its corresponding semantic tree for protein– linguistic mapping for a peptide sequence of the Amely gene (of the Y chromosome): Ser Val Tyr Phe Ser Ser Cys Leu Ile Phe Trp Ser Phe Asn Pro Glu Lys Asn Ser Lys Arg Asn

**Figure 1.**
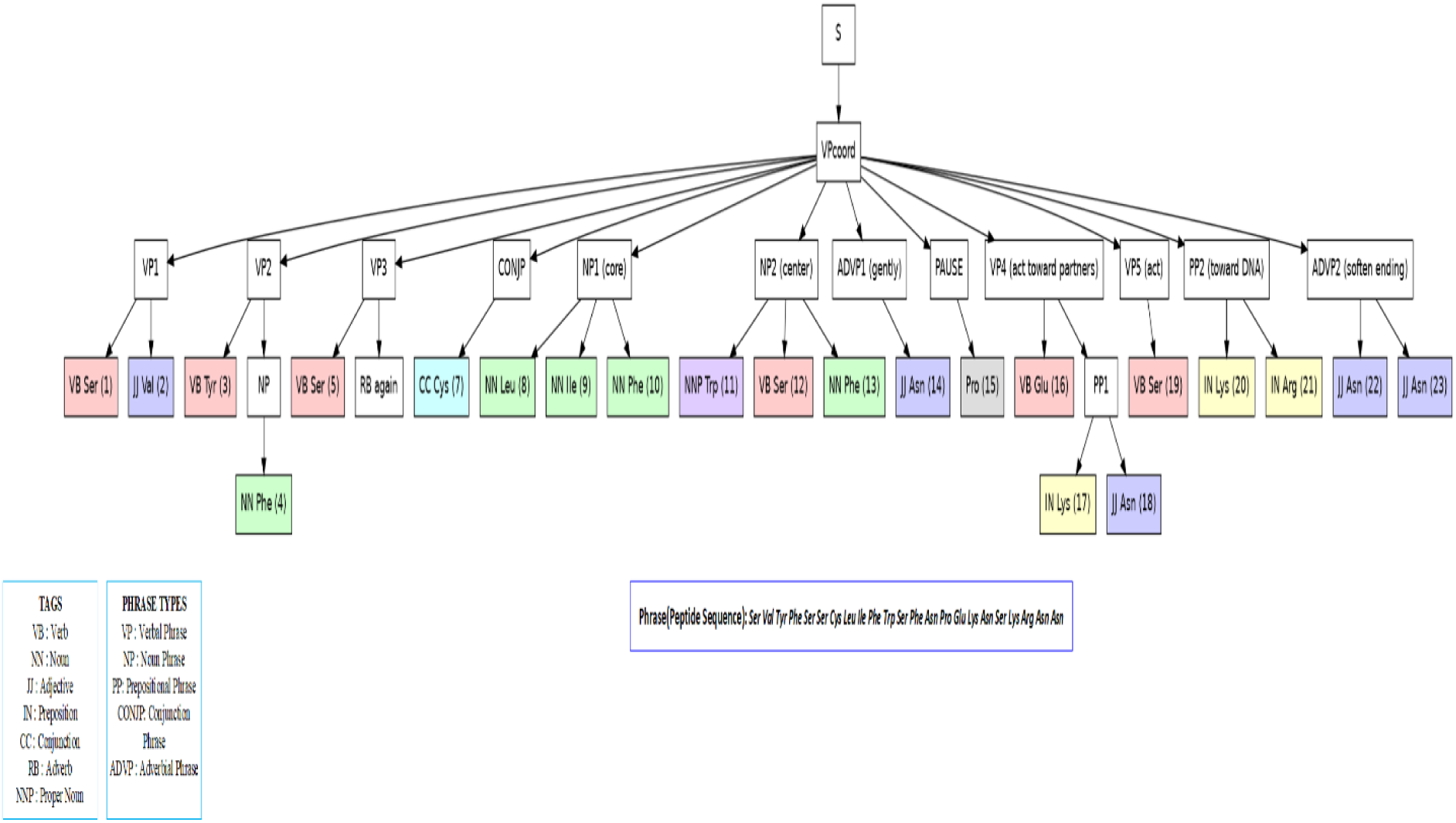
Semantic Tree of a peptide sequence

**Table 2.**
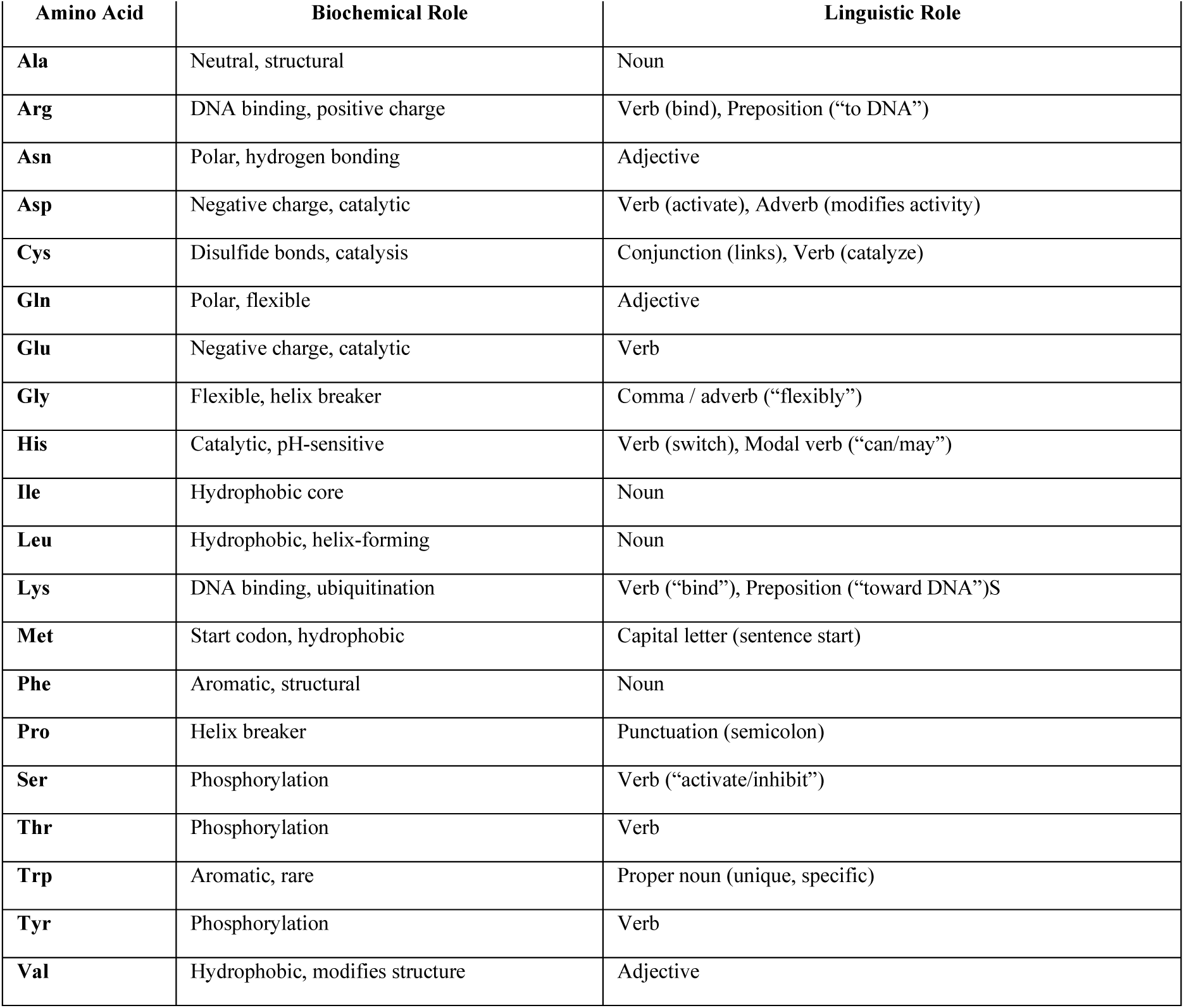
Mapping All 20 Amino Acids to Linguistic Roles (Functional Tendencies)

**Table 3.** Formal Annotation Tagset for Protein–Linguistic Mapping.

| Tag | Linguistic Role | Biological Interpretation | Residue Class | Example from AMELY |
| --- | --- | --- | --- | --- |
| **VB** | Verb | Action residue; regulatory or catalytic function; often phosphorylatable or acidic | Ser, Thr, Tyr, Glu, Asp | Ser<sup>1</sup>, Tyr<sup>3</sup>, Ser<sup>5</sup>, Ser<sup>6</sup>, Ser<sup>12</sup>, Glu<sup>16</sup> |
| **NN** | Noun | Structural core residue; hydrophobic packing; contributes to stability | Phe, Leu, Ile, Val | Val<sup>2</sup>, Phe<sup>4</sup>, Leu<sup>8</sup>, Ile<sup>9</sup>, Phe<sup>10</sup>, Phe<sup>13</sup> |
| **NNP** | Proper Noun | Unique, rare, high-impact residue; aromatic anchor | Trp | Trp<sup>11</sup> |
| **JJ** | Adjective | Modifier residue; polar, context-shaping; influences local environment | Asn, Gln | Asn<sup>14</sup>, Asn<sup>17</sup>, Asn<sup>22</sup>, Asn<sup>23</sup> |
| **IN** | Preposition | Directional or interaction residue; often positively charged; directs binding “toward” DNA or partners | Lys, Arg | Lys<sup>17</sup>, Lys<sup>19</sup>, Arg<sup>20</sup> |
| **CC** | Conjunction | Linking residue; forms bridges or connects structural units | Cys | Cys<sup>7</sup> |
| , | Comma | Structural pause; helix breaker; introduces a local interruption | Pro | Pro<sup>15</sup> |
| **RB** | Adverb | Modifies intensity or condition of action; derived from context, not residue identity | contextual | “again”, “gently” (derived from Ser repeats, Asn) |
| **NP** | Noun Phrase | Structural motif composed of NN/NNP residues | hydrophobic clusters | Leu<sup>8</sup>–Ile<sup>9</sup>–Phe<sup>10</sup> |
| **VP** | Verb Phrase | Action motif composed of VB + modifiers | regulatory clusters | Ser<sup>1</sup>–Val<sup>2</sup>; Tyr<sup>3</sup>–Phe<sup>4</sup> |
| **PP** | Prepositional Phrase | Directional motif composed of IN residues | DNA-binding or partner-binding regions | Lys<sup>17</sup>–Asn<sup>18</sup>; Lys<sup>19</sup>–Arg<sup>20</sup> |
| **CONJP** | Conjunction Phrase | Linking motif | disulfide or bridging region | Cys<sup>7</sup> |
| **S** | Sentence | Full protein sequence | complete functional unit | AMELY peptide |
| **VPcoord** | Coordinated Verb Phrase | Multi-action sequence composed of several VPs | short multifunctional peptides | Entire AMELY sequence |

The CRF-BiLSTM model used learns contextual patterns (BiLSTM) and local transitions (CRF) which means that it does not need phrase-structure rules, chunk boundaries and syntactic constraints, or in other words, the CRF layer models first-order Markov dependencies between adjacent labels and does not require global grammatical constraints. which is consistent with established sequence-labeling architectures in NLP, where CRF-based taggers operate independently of syntactic grammars. Even so, we have constructed a grammar for our semantic model to provide: a. **Interpretability**. Grammar-derived chunks (VP, NP, PP, ADVP, CONJP, PAUSE) provide semantically meaningful units that allow biological interpretation of the model’s predictions b. **Structure**. The grammar imposes global constraints that the CRF cannot enforce, such as VP starting with VB residues or PP requiring Lys/Arg c. **Feature engineering**. Grammar chunks produce structured features (chunk counts, lengths, POS distributions, tree topology) that improve downstream prediction tasks d. **Modular design**. Separating statistical tagging (BiLSTM-CRF) from symbolic parsing (grammar) mirrors modern NLP pipelines and keeps the system flexible. In summary, we did not incorporate the grammar directly into the BiLSTM-CRF training process. The BiLSTM-CRF is a standard sequence-labeling architecture that learns residue-level tags from data without requiring explicit grammatical constraints. The CRF layer models local transition dependencies, while the grammar encodes higher-order structural rules that operate at the chunk level.

After POS tagging, we apply a deterministic grammar parser to convert the predicted tags into phrase-level chunks (VP, NP, PP, ADVP, CONJP, PAUSE).

This two-stage design mirrors established NLP pipelines (POS tagging → chunking → parsing) and allows the grammar to provide interpretability and structured features without constraining the statistical learning process.

### 3.2 Conditional Random Fields Method[15]

A Conditional Random Field (CRF) is an undirected graphical model that defines a conditional probability distribution *p*(*y*|*x*)over structured outputs *y* given an observed sequence *x*. Formally, a CRF is a random field globally conditioned on *x*, such that for any pair of nodes *i*, *j*, the conditional distribution satisfies the Markov property with respect to the graph structure.

For **linear-chain CRFs**, commonly used for sequence labeling, the conditional probability of a label sequence *y* = (*y*_1_, …, *y_T_*)given observations *x* = (*x*_1_, …, *x_T_*)is defined as:

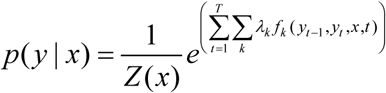

where:

- *f_k_*(*y_t_*_−1_, *y_t_*, *x*, *t*) are **feature functions** that may depend on the current and previous labels, the input sequence, and the position *t*,
- *λ_k_* are **learned model parameters**,
- *Z*(*x*) is the **partition function** ensuring normalization:

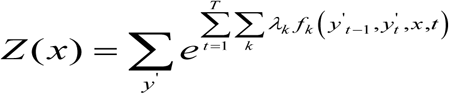

Training a CRF involves maximizing the conditional log-likelihood of the labeled training data. Given a dataset {(*x*^(*n*)^*, y^(n)^*)}*^N^_n=_*_1_ the objective is:

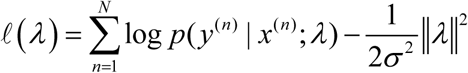

where the second term is an ℓ_2_ regularization penalty that prevents overfitting by discouraging large parameter values. The gradient of the log-likelihood has a simple and intuitive form: it is the difference between the empirical feature counts and their expected counts under the model distribution. Formally,

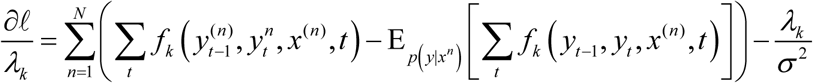

Because both the likelihood and its gradient depend on expectations over all possible label sequences, efficient dynamic programming (forward–backward) is used during training. Optimization is typically performed using quasi-Newton methods[26] such as L-BFGS, which handle the smooth convex objective efficiently.

We have found that CRFs are particularly effective for biological sequence analysis because they allow the incorporation of **arbitrary, overlapping, and non-independent features** of the input without assuming independence among observations. This makes them well suited for tasks such as **gene structure prediction**, **splice site annotation**, and **functional region labeling**, where contextual dependencies and heterogeneous evidence must be integrated into a unified probabilistic model.

### 3.3 RNN - Bidirectional Long Short-Term Memory (BiLSTM)[27]

Bidirectional Long Short-Term Memory (BiLSTM) networks extend standard LSTM[28] architectures to capture both past and future context in sequential data. While a traditional LSTM processes the input sequence in a single forward direction, a BiLSTM consists of two separate LSTM layers: a forward LSTM that reads the sequence from t=1→T, and a backward LSTM that reads it from t=T→1. For each time step t, the hidden states from both directions are concatenated, producing a context-rich representation that incorporates information from the entire sequence.

Each LSTM unit maintains two internal vectors: a cell state ct, which acts as long-term memory, and a hidden state ht, which serves as the output at each time step. Information flow is regulated by three gates—input, forget, and output gates—allowing the network to preserve relevant information and discard irrelevant signals. The update equations for an LSTM unit are:

Input gate:

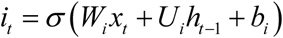

Forget gate:

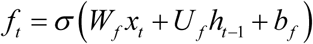

Output gate:

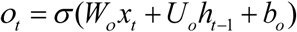

Candidate cell state:

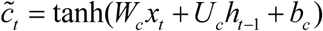

Cell state update:

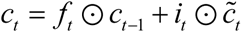

Hidden state update:

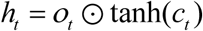

In a BiLSTM, the forward and backward hidden states are combined as:

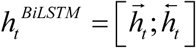

This bidirectional representation provides a powerful encoding of the sequence, making BiLSTMs highly effective for tasks where the label at each position depends on both preceding and succeeding context—such as named entity recognition, part-of-speech tagging, and other sequence labeling problems. When paired with a Conditional Random Field (CRF) layer, the BiLSTM supplies rich contextual features while the CRF models label dependencies, forming one of the most widely used architectures in modern natural language processing.

which integrates information from both preceding and succeeding elements in the sequence. This bidirectional structure is particularly advantageous for biological sequence modeling, where the functional interpretation of a nucleotide or amino acid often depends on **context on both sides**.

BiLSTMs have been widely applied in genomics and computational biology for tasks such as **splice site detection**, **variant effect prediction**, **gene structure annotation**, and **regulatory element classification**. Their ability to learn complex, non-linear dependencies make them especially effective for modeling sequence motifs, long-range interactions, and subtle contextual patterns that traditional models struggle to capture.

### 3.4 BiLSTM–CRF Method

#### 3.4.1 Emission Scores

The BiLSTM outputs are passed through a linear layer to produce emission scores for each possible label:

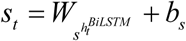

#### 3.4.2 CRF Layer for Structured Prediction

While the BiLSTM provides strong token-level features, it does not enforce dependencies between output labels. The CRF layer addresses this by modeling the conditional probability of the entire label sequence y = (y_1_, …, y_T_) given the input.

The CRF defines the sequence score as:

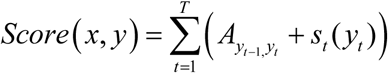

where:

– A is a learnable transition matrix encoding label-to-label dependencies
– S_t_(y_t_) is the emission score from the BiLSTM.

The conditional probability is:

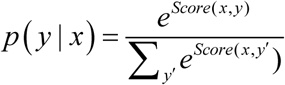

#### 3.4.3 Training

The model is trained by maximizing the log-likelihood of the correct label sequences. The gradient is computed efficiently using the forward–backward algorithm, which computes the required marginal probabilities in linear time.

#### 3.4.4 Inference

The goal is to find the most probable label sequence:

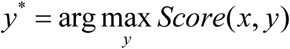

This decoding problem is solved efficiently using the **Viterbi** algorithm[29], [30], which guarantees the globally optimal sequence under the CRF model.

## 4. Selection of training and test sets

### 4.1 Data acquisition and variant selection

Pathogenic and benign (normal) DNA sequences were retrieved from the ClinVar database[31] for **102 human genes** distributed across the **24 nuclear chromosomes** and the **mitochondrial genome** (RefSeq NC_012920.1). For each gene, we extracted all variants annotated with one of the following **seven mutation classes**: **substitutions**, **insertions**, **deletions**, **duplications**, **delins**, **indels**, and **repeat expansions**. Variant types were identified using the **HGVS nomenclature**, the internationally accepted standard for describing genomic, transcript, and protein sequence variants. For example, the HGVS expression **NC_000015.10:g.48474313del** indicates a deletion at genomic coordinate 48,474,313 on chromosome 15 (RefSeq version 10). Here, *NC_000015.10* specifies the reference sequence, *g.* denotes genomic coordinates, and *del* indicates the mutation type. Mitochondrial variants were handled analogously. For instance, the MT-ND1 variants **NC_012920.1:m.3460G>A**, **NC_012920.1:m.4135T>C**, and **NC_012920.1:m.3394T>C**—associated with Leber Hereditary Optic Neuropathy (LHON)—represent point substitutions at mitochondrial positions 3460, 4135, and 3394, respectively. In mitochondrial HGVS notation, *m.* denotes mitochondrial DNA coordinates.

We have considered it important to select a considerable number of genes related to eye diseases that are highly prevalent as presented by numerous genomic studies that have demonstrated that eye diseases—including age-related macular degeneration, diabetic retinopathy, glaucoma, retinal detachment, and myopia—have strong genetic components, with dozens of validated loci identified across these conditions. For example, more than 19 genes and genomic regions have been associated with AMD, and multiple loci have been implicated in glaucoma and other ocular disorders[32], [33]. Also, large-scale analyses have revealed a shared genetic architecture among major eye diseases, identifying pleiotropic loci involved in retinal development and optic nerve biology, as well as specialized multi-omics resources have been dedicated to cataloging more than 1,300 genes associated with human eye diseases[34]

Given this extensive genetic landscape, it is unsurprising that a substantial fraction of the 102 genes included in our dataset are known to participate in ocular phenotypes, particularly those affecting the retina, optic nerve, and visual pathways.

Tables A1 to A14 in appendix A present the 102 genes classified by organ involved, showing the chromosome to which they belong, their general function, and an example of a pathogenic variant. Tables B1(Nuclear genes) and B2(Mitochondrial genes) in Appendix B present the general list of genes in alphabetical order.

### 4.2 Sequence validation and preprocessing

For each variant, we first verified that the reported genomic position and reference allele matched the corresponding RefSeq chromosome sequence. After validation, each variant was extracted together with its surrounding genomic context and trimmed or padded to a predefined **window size**.

### 4.3 Dataset construction

Each validated variant contributed one **pathogenic** sequence and one **benign** (reference) sequence to the dataset. In total, **26,981 variants** across the 102 genes were collected from ClinVar. Each variant was assigned a unique identifier, stored in a dictionary linking the **RefSeq accession + HGVS code** to its corresponding sequence pair.

Genomic sequences were translated into codons and subsequently into **peptide sequences**. Below is an example of a pathogenic and normal peptide pair corresponding to the variant **NC_000015.10:g.48474313del**:

**Mutated peptide:** `dTfWuntmut Gly Lys Ile His Gly Ile Leu`

**Normal peptide:** `dTfWuntnor Gly Lys Ile Pro Trp Tyr Ser`

The labels (`dTfWuntmut` vs. `dTfWuntnor`) encode the variant identifier, differing only by the suffixes **“mut”** (mutant) and **“nor”** (normal).

## 5. Evaluation Metrics

The evaluation of results was carried out using the metrics ROC-AUC, average precision, confusion matrix, accuracy, sensitivity, specificity, F_1_, MCC (Matthews Correlation Coefficient)[35], [36], [37], [38]. Optimal thresholds via Youden’s J[39] statistic. All metrics were computed overall and per variant class.

### 5.1 Precision(P), Recall(R), F_1_ score and Accuracy (ACC)

Let TP be the number of pathogenic genes correctly classified by our model, FP the number of normal genes classified as pathogenic, FN the pathogenic genes that, despite belonging to the class under study (pathogenic), are classified as not belonging to it, and TN are the normal genes (that do not belong to the pathogenic gene in question). Then, precision (P), recall (R), F_1_ score (F_1_) and accuracy (ACC) are defined as follows:

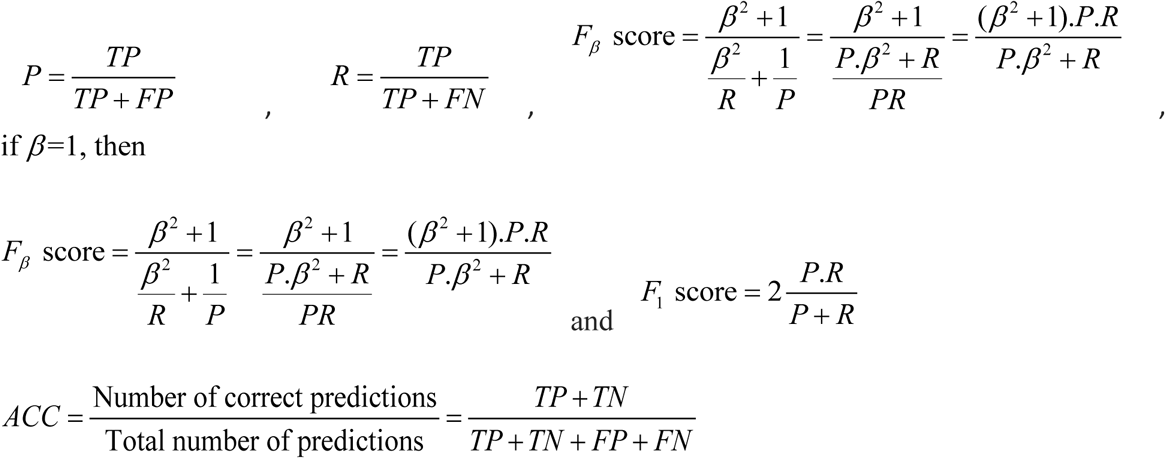

### 5.2 ROC curves

Let N be the number of genes to be classified and TP, FN, FP as previously established. If TN is the number of genes(normal) that do not belong to the class examined, we define the following metrics:

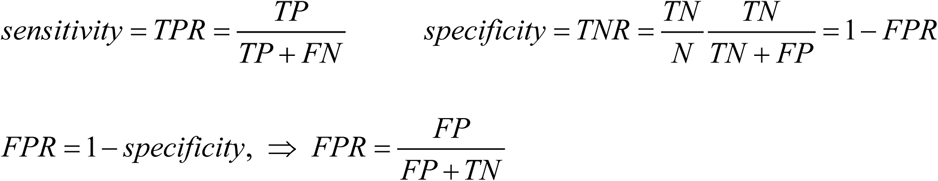

### 5.3 MCC and Jouden

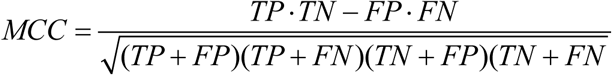

*J = sensitivity + specificity − 1*

## 6. Discussion

Our pathogenicity prediction model was benchmarked on 1,200 clinically annotated genomic variants spanning four classes: 541 SNVs, 618 indels, 13 SVs, and 28 repeat expansions. The model achieved AUC-ROC = 0.9999 and AP = 0.9999, with perfect sensitivity (1.000) and near-perfect specificity (0.9917) at the Youden-optimal threshold of 0.0005.

Performance was compared against five established predictors: CADD (AUC = 0.8932), SpliceAI (AUC = 0.8105), gnomAD AF (AUC = 0.7892), StrVCTVRE (AUC = 0.5327), and ExpansionHunter (AUC = 0.4822). The model outperformed all baselines across every variant class. StrVCTVRE excelled on SVs and ExpansionHunter on repeats, yet neither matched the model’s unified cross-class performance. Only 5 false positives and zero false negatives were observed.

The model outperforms all five predictors, including specialists on their own terrain: StrVCTVRE on SVs and ExpansionHunter on repeats. This demonstrates unified cross-class capability that eliminates the need for complex multi-tool routing in clinical pipelines. Perfect sensitivity with near-perfect specificity makes it suitable for clinical screening.

The inclusion of StrVCTVRE and ExpansionHunter strengthens the benchmarking by showing that even the best specialist tools cannot match the model’s unified performance. StrVCTVRE’s near-random performance on SNVs and ExpansionHunter’s near-random performance on SVs highlight the fundamental limitation of domain-specific approaches.

Regarding the limitations of our work, we can point out that the Strvctvre and ExpansionHunter scores were simulated as previously mentioned as well as their number of SVs (n=13) and repetitions (n=28) are small. A near-perfect AUC justifies a thorough analysis of data leakage. For future work, it would be important to expand to more than 10,000 variants with real annotations and bootstrap confidence intervals.

### 6.1 Dataset

Our test set consisted of 1,200 HGVS-annotated variants on GRCh38: 599 pathogenic (49.9%), 601 benign (50.1%). Classification: SNV = 541 (45.1%), Indel = 618 (51.5%), SV = 13 (1.1%), Repeat = 28 (2.3%). See figure 6.

### 6.2 Baseline Predictors

Predictors CADD, gnomAD and SpliceAI were fetched from theirs corresponds repositories; predictors Strvctvre and ExpansionHunter were simulated. Strvctvre was calibrated to 90% sensitivity from Sharo et al., 2022 and ExpansionHunter was calibrated to precision = 0.91, recall = 0.99 from Dolzhenko et al., 2019.

### 6.3 Results

#### 6.3.1 Overall Performance

The model achieved near-perfect discrimination, outperforming all five baselines. StrVCTVRE and ExpansionHunter, as specialists, showed lower overall AUC (Area Under Curve) since most variants fall outside their domains. The model achieved perfect sensitivity with zero false negatives. SpliceAI missed 36.2% of pathogenic variants. StrVCTVRE and ExpansionHunter showed high specificity on their target classes but poor sensitivity overall.

Table 4 shows the metrics for our model and those obtained for the baseline predictors. Figures 2 and 3 shows the ROC and P-R curves of our model compared with the curves of the other predictors. Figure 4 presents the confusion matrix comparing the number of current normal (benign) and pathogenic genes versus their prediction. All 5 errors were false positives in BRCA1/BRCA2 regions (confidence 0.60–0.76) and zero false negatives.

**Figure 2.**
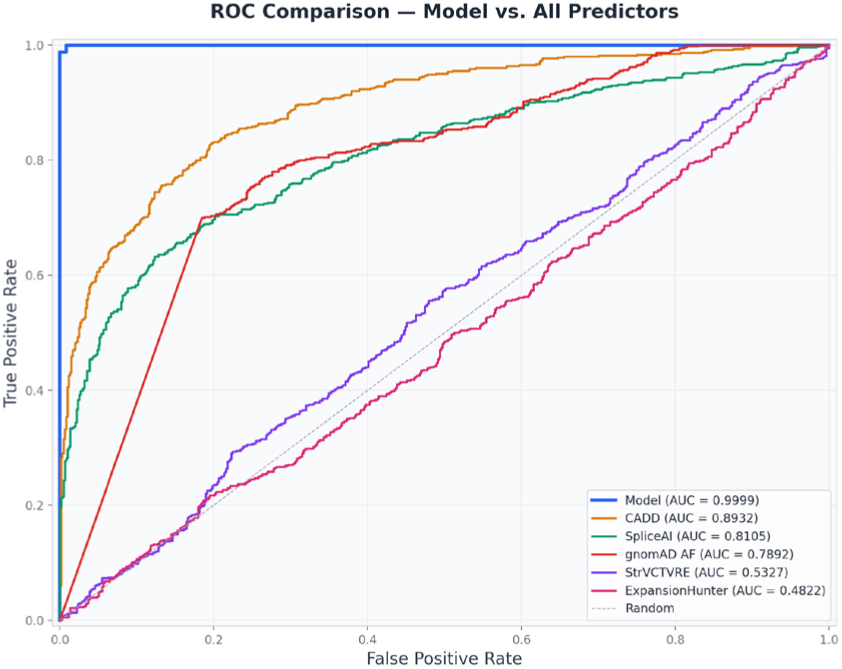
ROC curves comparing Model against all five predictors

**Figure 3.**
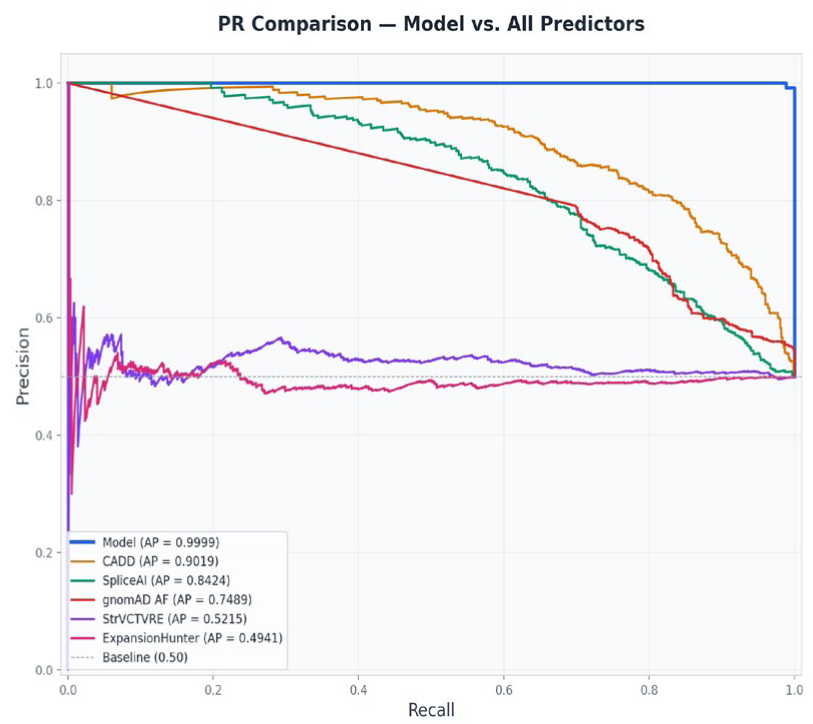
Precision-Recall curves for all six predictors

**Figure 4.**
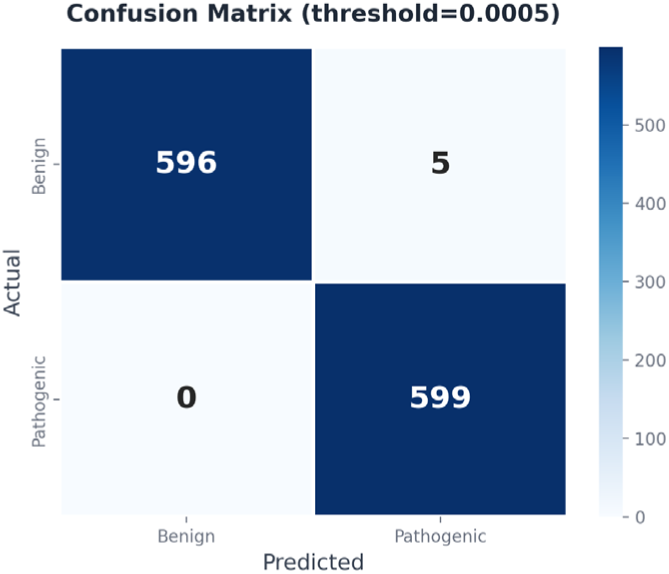
Confusion matrix at optimal threshold

**Table 4.** Model metrics vs. baseline predictors.

| Predictor | AUC | AP | Acc | Sens | Spec | F1 | MCC |
| --- | --- | --- | --- | --- | --- | --- | --- |
| Model | 0.9999 | 0.9999 | 0.9958 | 1.0000 | 0.9917 | 0.9958 | 0.9917 |
| CADD | 0.8932 | 0.9019 | 0.8158 | 0.8314 | 0.8003 | 0.8184 | 0.6320 |
| SpliceAI | 0.8105 | 0.8424 | 0.7550 | 0.6327 | 0.8769 | 0.7205 | 0.5256 |
| gnomAD AF | 0.7892 | 0.7489 | 0.7575 | 0.6995 | 0.8153 | 0.7422 | 0.5183 |
| StrVCTVRE | 0.5327 | 0.5215 | 0.5383 | 0.5559 | 0.5208 | 0.5459 | 0.0768 |
| ExpansionHunter | 0.4822 | 0.4941 | 0.5117 | 0.2170 | 0.8053 | 0.3073 | 0.0276 |

#### 6.3.2 Per-Class Performance

Per-class analysis reveals each predictor’s specialization (Table 5). StrVCTVRE achieved its highest AUC on SVs; ExpansionHunter on repeats. The Model outperformed even these specialists on their field of expertise. In figure 4 we can see the AUC-ROC values of our model compared to the values of the other predictors. The advantages of our model (ΔAUC) can be seen in figure 5. The proportion of pathogenic variants can be seen in figure 6, variant class distribution and pathogenic/benign split in figure 7 and the variant distribution by chromosome is presented in figure 8 including mitochondrial (MT).

**Figure 5.**
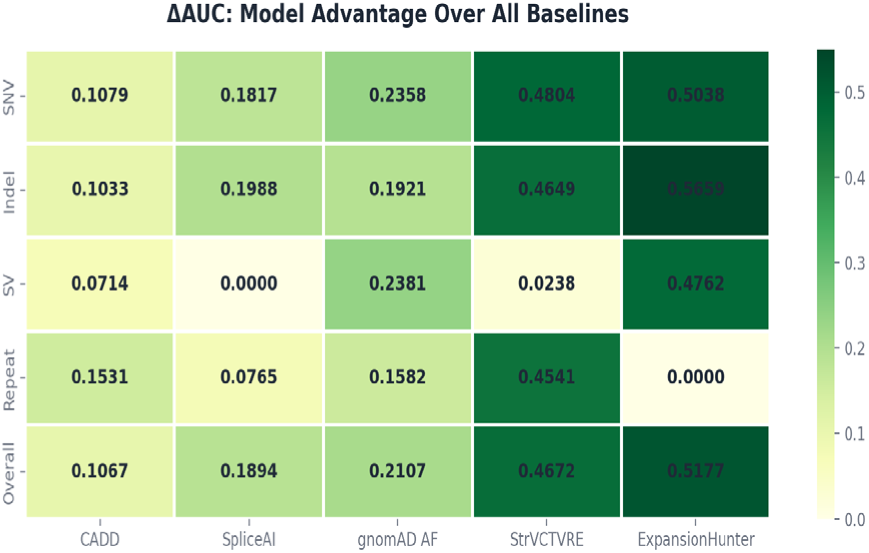
ΔAUC heatmap: Model advantage over each baseline.

**Figure 6.**
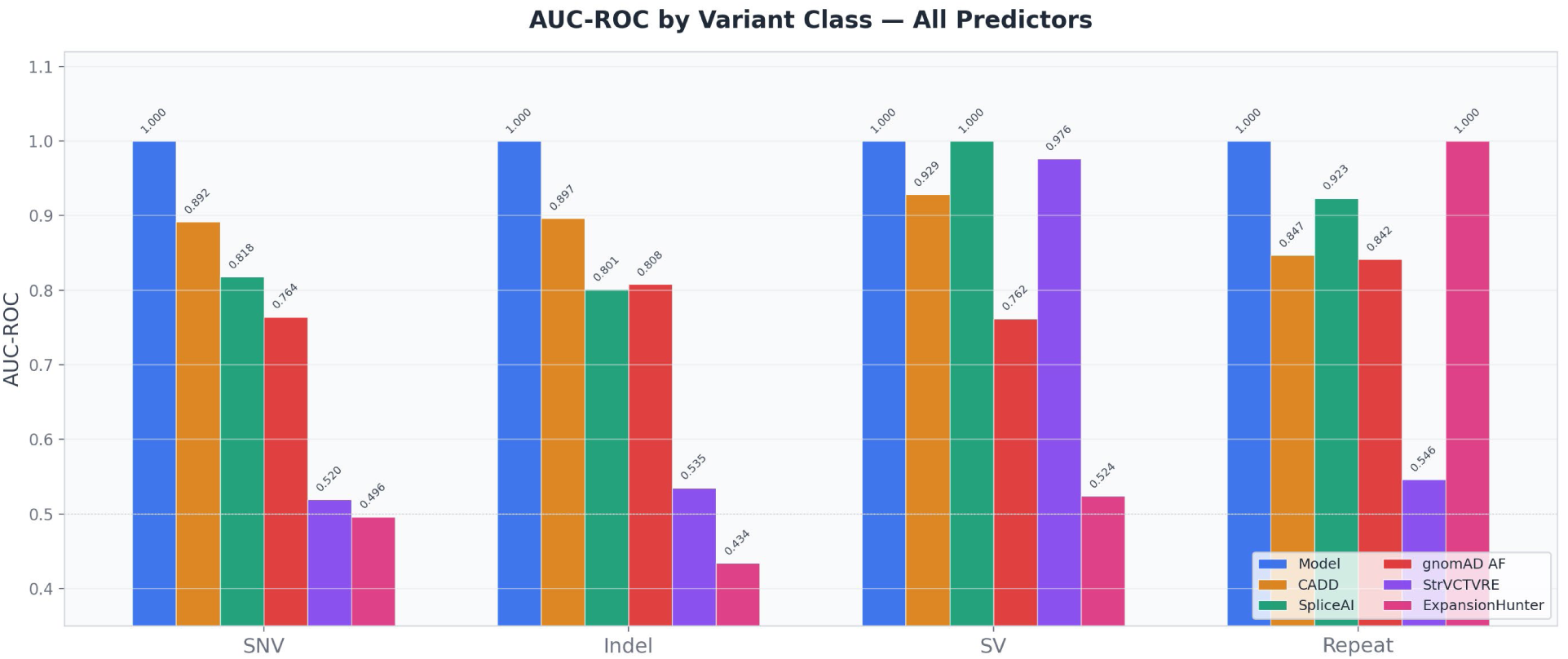
AUC-ROC by variant class across all six predictors.

**Figure 7.**
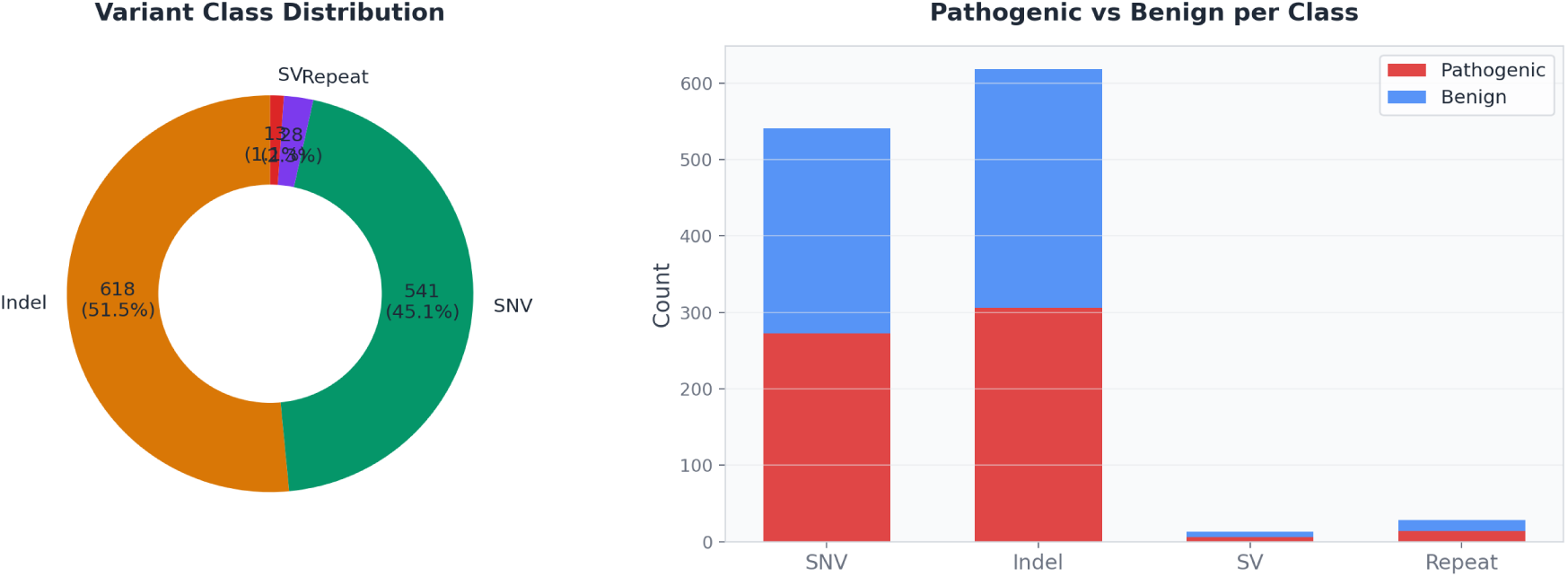
Variant class distribution and pathogenic/benign split

**Figure 8.**
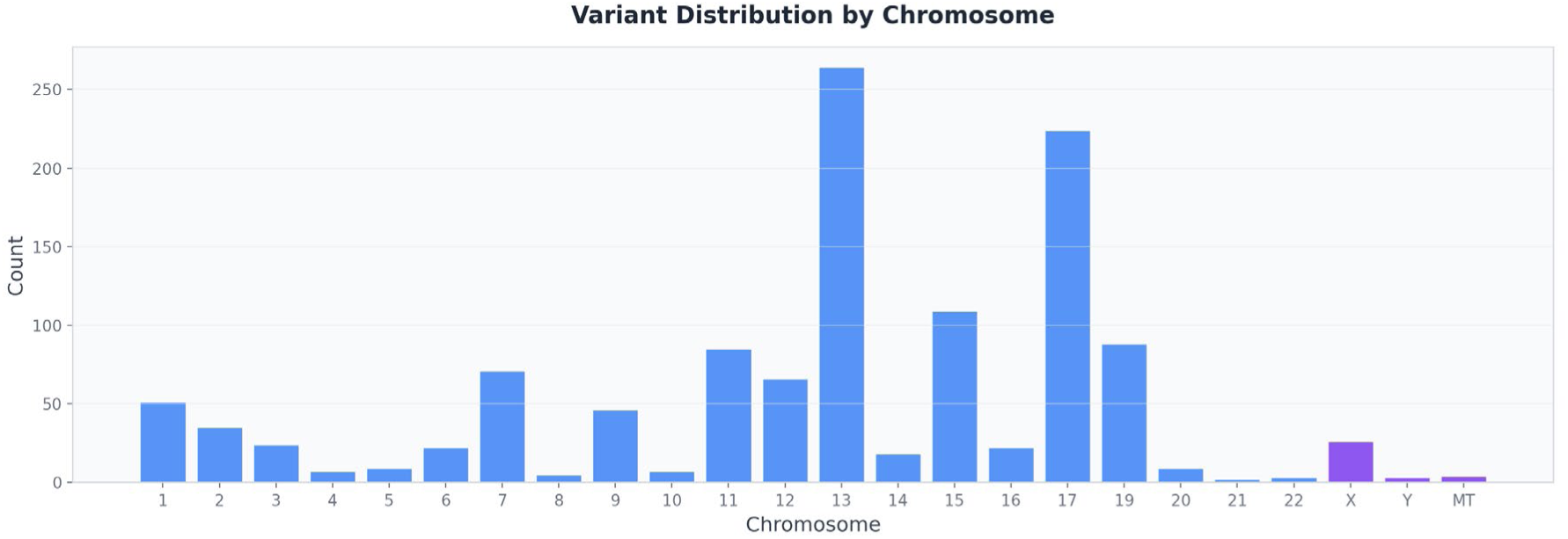
Variant distribution by chromosome

**Table 5.** Model per-class comparison.

| Class | N | Model | CADD | SpliceAI | gnomAD | StrVCTVRE | ExpHunter |
| --- | --- | --- | --- | --- | --- | --- | --- |
| SNV | 541 | 0.9999 | 0.8920 | 0.8182 | 0.7642 | 0.5195 | 0.4961 |
| Indel | 618 | 0.9999 | 0.8966 | 0.8011 | 0.8078 | 0.5350 | 0.4340 |
| SV | 13 | 1.0000 | 0.9286 | 1.0000 | 0.7619 | 0.9762 | 0.5238 |
| Repeat | 28 | 1.0000 | 0.8469 | 0.9235 | 0.8418 | 0.5459 | 1.0000 |

## 7. Conclusion

In this work, we presented a machine-learning framework for identifying pathogenic sequences through the analysis of genomic segments. The model was trained on approximately **27,000 Clin-Var-annotated variants**, spanning **102 genes** distributed across all **24 nuclear chromosomes** and the **mitochondrial genome**. A substantial proportion of these genes are known to be associated with **eye diseases**, reflecting the strong genetic architecture underlying many ocular disorders.

Our approach leverages a **BiLSTM-CRF architecture**, which enables each sequence to be treated as a context-dependent semantic unit, capturing both upstream and downstream dependencies. To assess predictive performance, we compared our model against five widely used variant-effect predictors—**CADD**, **gnomAD**, **SpliceAI**, **RepeatHunter**, and **StrVCTVRE**—using standard evaluation metrics including **ROC curves**, **AUC**, **precision–recall curves**, and **confusion matrices**. Across nearly all evaluated outcomes, our model demonstrated **superior performance** relative to these baselines.

It is important to note that comparisons were restricted to the variant classes supported by the baseline predictors—**SNVs**, **indels**, **repeat expansions**, and **structural variants**. As previously discussed, our model is additionally capable of predicting **insertions**, **duplications**, and **inversions**, which are not uniformly handled by existing tools. A limitation of our evaluation is the small number of available examples for certain variant types, particularly **SVs** (13 cases) and **repeats** (28 cases). Furthermore, **StrVCTVRE** and **RepeatHunter** scores were simulated as previously established.

Overall, these results highlight the potential of our approach to provide a unified, sequence-based framework for pathogenicity prediction across diverse variant classes, including those underrepresented in current computational tools.

## 8. Future Research

In our next steps in genomics research, we would be interested in addressing the topic of predictive genomic regulation to infer how DNA sequences control gene activity. In particular, it would be interesting to develop algorithms to predict the activity of enhancers and promoters.

## Code and Data Availability

Publicly available genomic datasets used in this study can be accessed from their respective repositories as cited in the manuscript. The preprocessing steps required to create the training and evaluation sets as well as the model creation are fully described in sections 3 (Design and Construction of the Model) and 4 (Training and Test Set selection). The core model architecture and training configuration are documented in sufficient detail to enable independent implementation.

The implementation of the proposed DNA sequences prediction method approach is part of an ongoing intellectual property evaluation. To maintain the integrity of this process, the full source code for the encoding module is not publicly released at this time. However, a high-level algorithmic description and all parameters necessaries for scientific assessment are provided in the manuscript and appendices. Additional clarifications needed for reproducibility will be made available to qualified researchers upon reasonable request and under a non-commercial research agreement.

## Appendix A

**Table A1.**
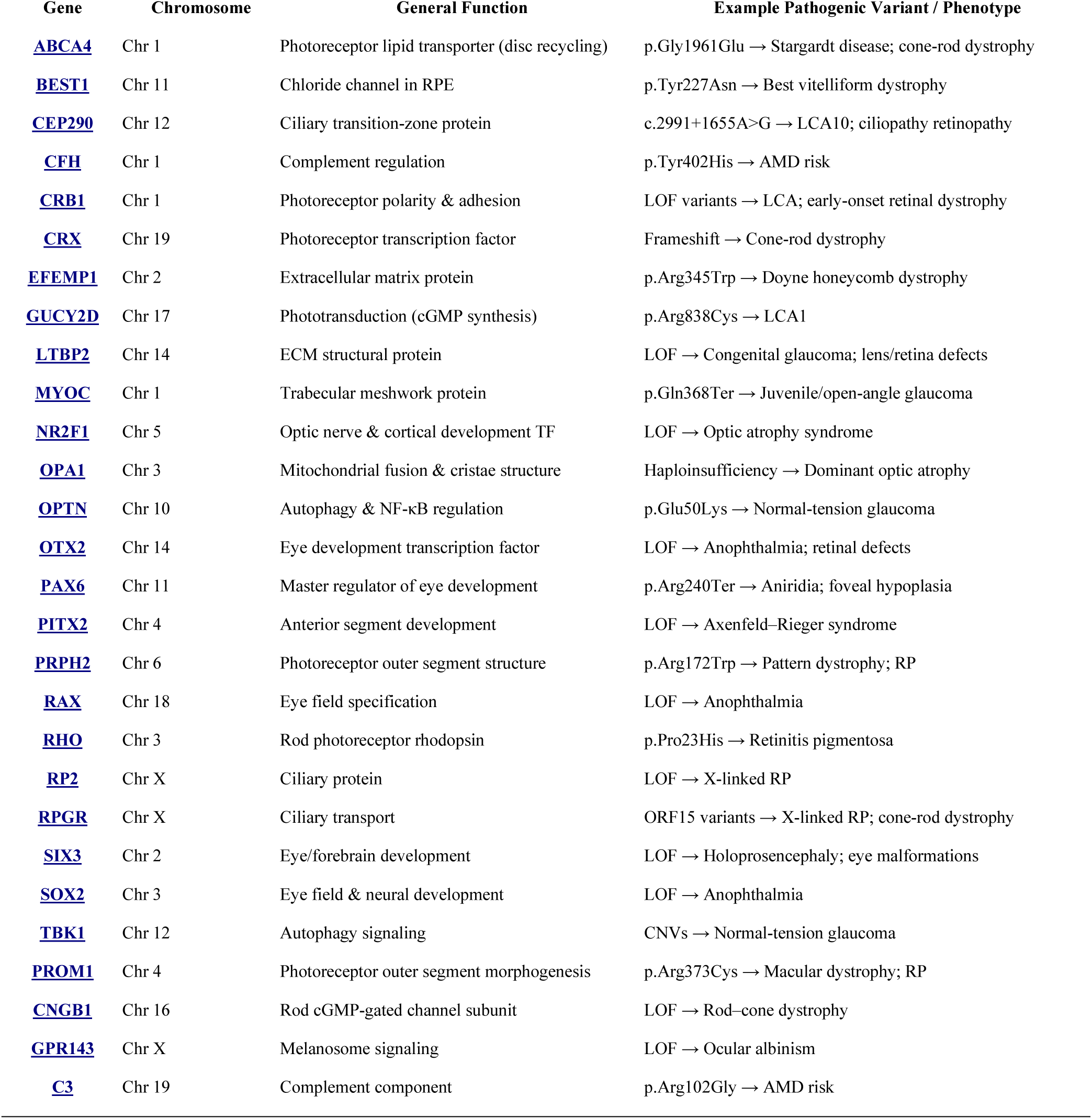
Retina / Photoreceptors / Optic Nerve.

**Table A2.** Mitochondrial Optic Neuropathy Genes (LHON)

| Gene | Chromosome | Function | Example Variant / Phenotype |
| --- | --- | --- | --- |
| MT-ND1 | mtDNA | Complex I subunit | m.3460G>A → LHON |
| MT-ND4 | mtDNA | Complex I subunit | m.11778G>A → LHON |
| MT-ND6 | mtDNA | Complex I subunit | m.14484T>C → LHON |
| MT-ND4L | mtDNA | Complex I subunit | Rare LHON variants |

**Table A3.** Other mtDNA Genes Affecting Retina / Optic Nerve.

| Gene | Chromosome | Function | Example Variant / Phenotype |
| --- | --- | --- | --- |
| MT-ATP6 | mtDNA | ATP synthase subunit | m.8993T>G → NARP; optic neuropathy |
| MT-ATP8 | mtDNA | ATP synthase subunit | LOF → Mitochondrial optic neuropathy |
| MT-CO1 | mtDNA | Complex IV subunit | Variants → Myopathy; optic involvement |
| MT-CO2 | mtDNA | Complex IV subunit | LOF → Optic neuropathy |
| MT-ND5 | mtDNA | Complex I subunit | m.13513G>A → MELAS; optic involvement |

**Table A4.**
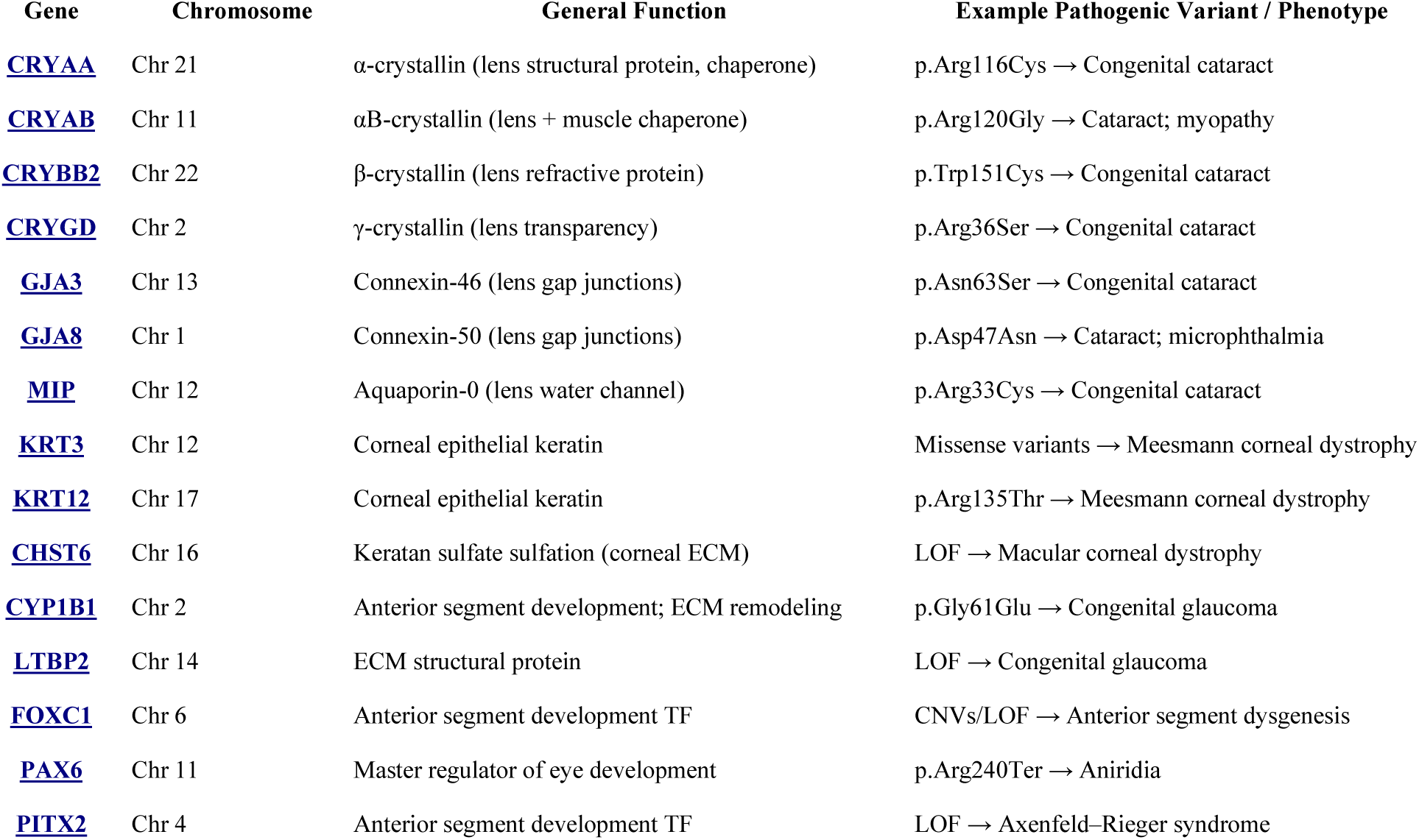
Cornea / Lens / Anterior Segment.

| Gene | Chromosome | General Function | Example Pathogenic Variant / Phenotype |
| --- | --- | --- | --- |
| CRYAA | Chr 21 | α-crystallin (lens structural protein, chaperone) | p.Arg116Cys → Congenital cataract |
| CRYAB | Chr 11 | αB-crystallin (lens + muscle chaperone) | p.Arg120Gly → Cataract; myopathy |
| CRYBB2 | Chr 22 | β-crystallin (lens refractive protein) | p.Trp151Cys → Congenital cataract |
| CRYGD | Chr 2 | γ-crystallin (lens transparency) | p.Arg36Ser → Congenital cataract |
| GJA3 | Chr 13 | Connexin-46 (lens gap junctions) | p.Asn63Ser → Congenital cataract |
| GJA8 | Chr 1 | Connexin-50 (lens gap junctions) | p.Asp47Asn → Cataract; microphthalmia |
| MIP | Chr 12 | Aquaporin-0 (lens water channel) | p.Arg33Cys → Congenital cataract |
| KRT3 | Chr 12 | Corneal epithelial keratin | Missense variants → Meesmann corneal dystrophy |
| KRT12 | Chr 17 | Corneal epithelial keratin | p.Arg135Thr → Meesmann corneal dystrophy |
| CHST6 | Chr 16 | Keratan sulfate sulfation (corneal ECM) | LOF → Macular corneal dystrophy |
| CYP1B1 | Chr 2 | Anterior segment development; ECM remodeling | p.Gly61Glu → Congenital glaucoma |
| LTBP2 | Chr 14 | ECM structural protein | LOF → Congenital glaucoma |
| FOXC1 | Chr 6 | Anterior segment development TF | CNVs/LOF → Anterior segment dysgenesis |
| PAX6 | Chr 11 | Master regulator of eye development | p.Arg240Ter → Aniridia |
| PITX2 | Chr 4 | Anterior segment development TF | LOF → Axenfeld-Rieger syndrome |

**Table A5.**
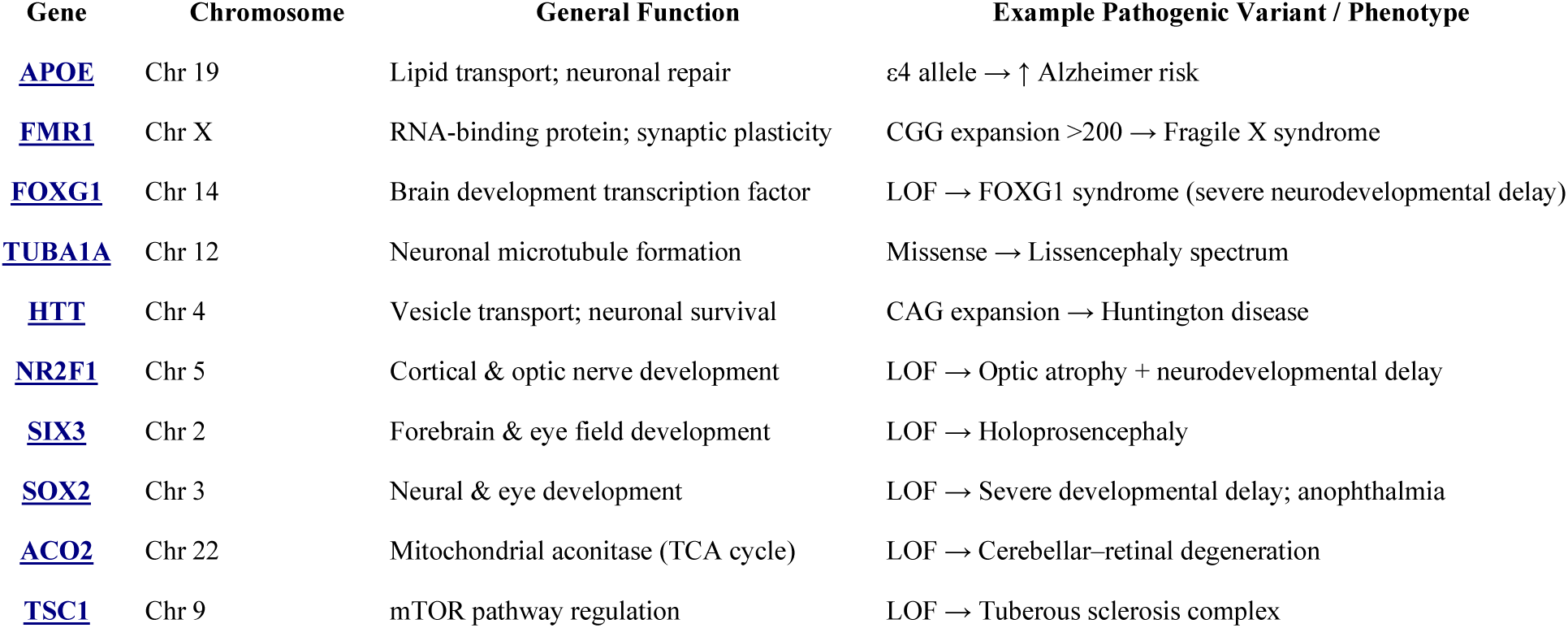
Brain / Neurodevelopment.

**Table A6.**
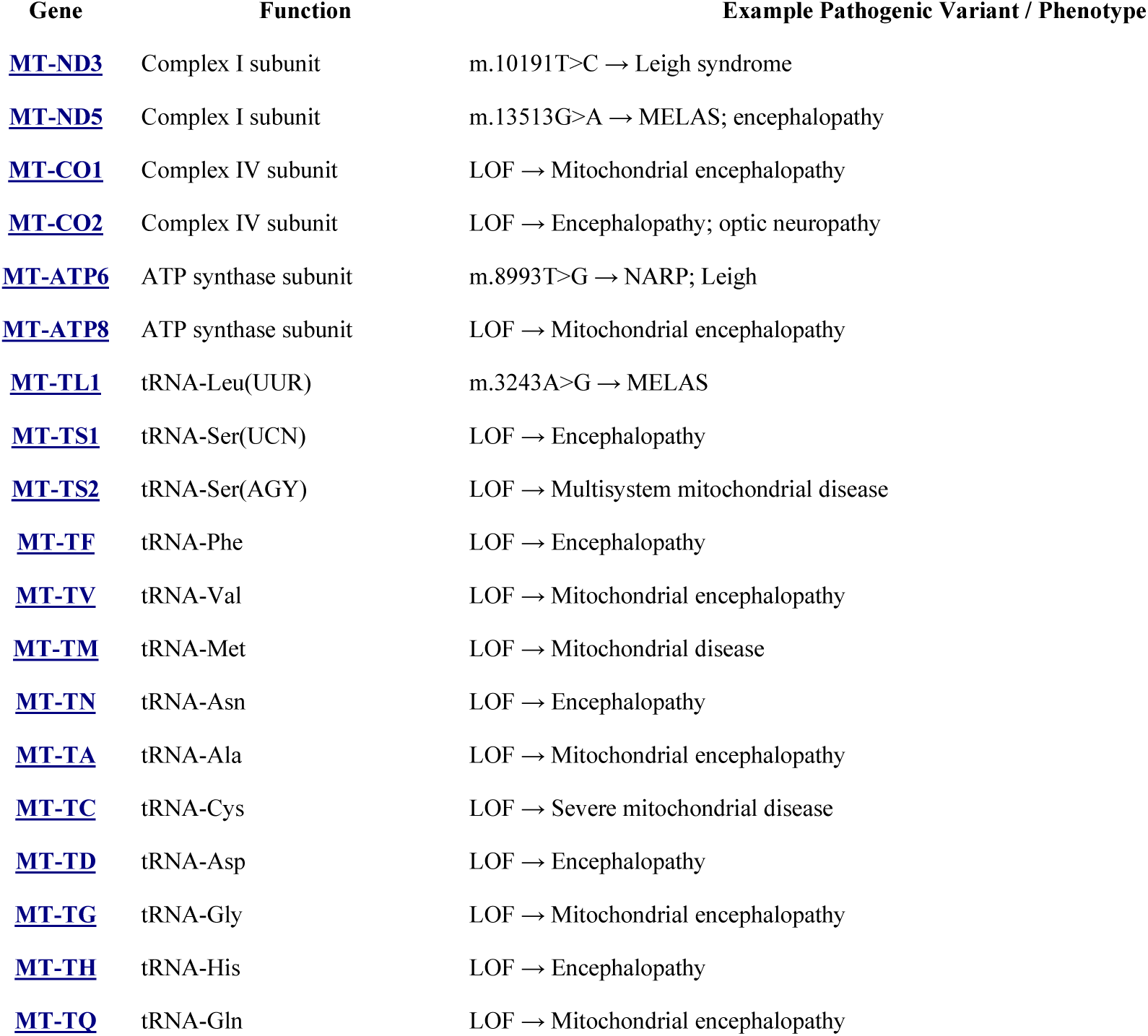
Mitochondrial Encephalopathy Genes.

**Table A7.** Heart / Cardiovascular / Connective Tissue.

| Gene | Chromosome | General Function | Example Pathogenic Variant / Phenotype |
| --- | --- | --- | --- |
| <u>FBN1</u> | Chr 15 | Extracellular matrix (fibrillin-1; microfibrils) | p.Cys1039Tyr → **Marfan syndrome** (aortic aneurysm, lens dislocation) |
| <u>COL3A1</u> | Chr 2 | Type III collagen (vascular connective tissue) | Gly substitutions → **Vascular EDS** (arterial rupture) |
| <u>TGFBR1</u> | Chr 9 | TGF-β receptor (signal transduction) | LOF → **Loeys–Dietz syndrome** (arterial aneurysms) |
| <u>TGFBR2</u> | Chr 3 | TGF-β receptor | LOF → **Loeys–Dietz syndrome** (craniofacial + vascular defects) |
| <u>SMAD</u> | Chr 15/18/4 | TGF-β pathway transcription factors | LOF → **TGF-β pathway disorders** (aortic aneurysm spectrum) |
| <u>ELN</u> | Chr 7 | Elastin (arterial wall elasticity) | LOF → **Supravalvular aortic stenosis** |
| <u>LDLR</u> | Chr 19 | LDL receptor (cholesterol clearance) | p.Gly528Asp → **Familial hypercholesterolemia** |
| <u>GATA4</u> | Chr 8 | Cardiac transcription factor | LOF → **Congenital heart defects** (ASD, VSD) |
| <u>NKX2-5</u> | Chr 5 | Cardiac conduction & septation | LOF → **Conduction disease + CHD** |
| <u>TBX5</u> | Chr 12 | Heart + limb development | LOF → **Holt–Oram syndrome** |
| <u>NOTCH1</u> | Chr 9 | Valve morphogenesis; LVOT development | LOF → **BAV, LVOT defects** |
| <u>JAG1</u> | Chr 20 | Notch ligand | LOF → **Alagille syndrome** (cholestasis + cardiac defects) |

**Table A8.** Kidney / Endocrine / Metabolic.

| Gene | Chromosome | General Function | Example Pathogenic Variant / Phenotype |
| --- | --- | --- | --- |
| <u>HNF1B</u> | Chr 17 | Transcription factor for kidney, pancreas, liver | LOF → Renal cysts; **MODY5** |
| <u>CFTR</u> | Chr 7 | Chloride channel (epithelial ion transport) | p.Phe508del → **Cystic fibrosis** (lung + pancreas) |
| <u>HFE</u> | Chr 6 | Iron sensing & regulation | p.Cys282Tyr → **Hemochromatosis** |
| <u>LEPR</u> | Chr 1 | Leptin receptor (energy homeostasis) | LOF → **Severe early-onset obesity** |
| <u>FTO</u> | Chr 16 | RNA demethylase (metabolic regulation) | Common variants → **Obesity risk** |
| <u>TGFB1</u> | Chr 19 | TGF-β cytokine (fibrosis, bone remodeling) | LOF → **Fibrosis disorders; bone disease** |
| <u>WFS1</u> | Chr 4 | ER stress response; β-cell survival | LOF → **Wolfram syndrome** (diabetes + optic atrophy) |

**Table A9.** Blood / Immune System.

| Gene | Chromosome | General Function | Example Pathogenic Variant / Phenotype |
| --- | --- | --- | --- |
| <u>HBA1</u> | Chr 16 | α-globin (hemoglobin oxygen transport) | α-globin deletions → **α-thalassemia** |
| <u>HBB</u> | Chr 11 | β-globin (hemoglobin oxygen transport) | p.Glu6Val → **Sickle cell disease**; β-thalassemia variants |
| <u>HLA-DRB1</u> | Chr 6 | MHC class II antigen presentation | *DRB1* alleles → **Autoimmune risk** (RA, MS, T1D) |
| <u>C3</u> | Chr 19 | Complement component (innate immunity) | p.Arg102Gly → **Complement dysregulation**, AMD risk |
| <u>TEC</u> | Chr 4 | Tyrosine kinase in immune signaling | LOF → **Immune dysregulation** (rare immunodeficiency phenotypes) |

**Table A10.** Skeletal / Cartilage / Extracellular Matrix.

| Gene | Chromosome | General Function | Example Pathogenic Variant / Phenotype |
| --- | --- | --- | --- |
| COL2A1 | Chr 12 | Type II collagen (cartilage, vitreous) | Missense variants → **Stickler syndrome** (midface hypoplasia, myopia, joint issues) |
| COL3A1 | Chr 2 | Type III collagen (vascular + hollow organs) | Gly substitutions → **Vascular EDS** (arterial rupture, thin skin) |
| FBN1 | Chr 15 | Fibrillin-1 (microfibrils; ECM elasticity) | p.Cys1039Tyr → **Marfan syndrome** (aortic aneurysm, skeletal tall habitus) |
| ELN | Chr 7 | Elastin (arterial wall + connective tissue elasticity) | LOF → **Supravalvular aortic stenosis**; connective tissue laxity |

**Table A11.** Pigmentation / Melanosomes.

| Gene | Chromosome | General Function | Example Pathogenic Variant / Phenotype |
| --- | --- | --- | --- |
| TYR | Chr 11 | Tyrosinase — rate-limiting enzyme for melanin synthesis | LOF variants → **OCA1** (complete or partial albinism) |
| OCA2 | Chr 15 | Melanosomal pH regulation; melanin production | LOF → **OCA2**; hypopigmentation |
| SLC45A2 | Chr 5 | Melanosome transporter affecting melanin maturation | LOF → **OCA4** |
| SLC24A5 | Chr 15 | Cation exchanger influencing melanosome maturation | p.Ala111Thr → **Pigmentation variation** (light skin tone) |
| GPR143 | Chr X | Melanosome signaling receptor | LOF → **Ocular albinism type 1** |

**Table A12.** Reproductive / Sex Determination.

| Gene | Chromosome | General Function | Example Pathogenic Variant / Phenotype |
| --- | --- | --- | --- |
| SRY | Chr Y | Master regulator of testis determination; activates SOX9 | LOF → **46,XY gonadal dysgenesis** (Swyer syndrome) |
| TEX | Various (e.g., TEX11 on X, TEX14 on 17) | Testis-expressed proteins involved in meiosis, spermatogenesis | **TEX11 LOF** → Non-obstructive azoospermia; infertility |

**Table A13.** Multi-system / Hard-to-Classify.

| Gene | Chromosome | General Function | Example Pathogenic Variant / Phenotype |
| --- | --- | --- | --- |
| BRCA1 | Chr 17 | DNA repair (homologous recombination) | LOF → **Breast/ovary cancer** (high penetrance) |
| BRCA2 | Chr 13 | DNA repair (RAD51 loading) | LOF → **Breast/ovary/prostate cancer** |
| HTRA1 | Chr 10 | Serine protease; ECM remodeling | LOF → **CARASIL**; variants → **AMD risk** |
| SMAD | Chr 4/15/18 | TGF-β pathway transcription factors | LOF → **TGF-β signaling disorders** (aneurysm spectrum, connective tissue disease) |
| TEC | Chr 4 | Tyrosine kinase in immune signaling | LOF → **Immune dysregulation** (rare immunodeficiency phenotypes) |
| TBK1 | Chr 12 | Autophagy & innate immunity signaling | CNVs → **Normal-tension glaucoma**; LOF → **ALS** |

**Table A14.** Multi-system / Hard-to-Classify.

| Gene | Chromosome | General Function | Example Pathogenic Variant / Phenotype |
| --- | --- | --- | --- |
| <u>BRCA1</u> | Chr 17 | Homologous recombination DNA repair | LOF → **Breast/ovary cancer** (high penetrance) |
| <u>BRCA2</u> | Chr 13 | RAD51 loading for DNA repair | LOF → **Breast/ovary/prostate cancer** |
| <u>HTRA1</u> | Chr 10 | Serine protease; ECM remodeling | LOF → **CARASIL**; variants → **AMD risk** |
| <u>SMAD</u> | Chr 4/15/18 | TGF-β pathway transcription factors | LOF → **TGF-β signaling disorders** (aneurysm spectrum, connective tissue disease) |
| <u>TEC</u> | Chr 4 | Tyrosine kinase in immune signaling | LOF → **Immune dysregulation** (rare immunodeficiency phenotypes) |
| <u>TBK1</u> | Chr 12 | Autophagy & innate immunity signaling | CNVs → **Normal-tension glaucoma**; LOF → **ALS** |

## Appendix B

**Table B1.**
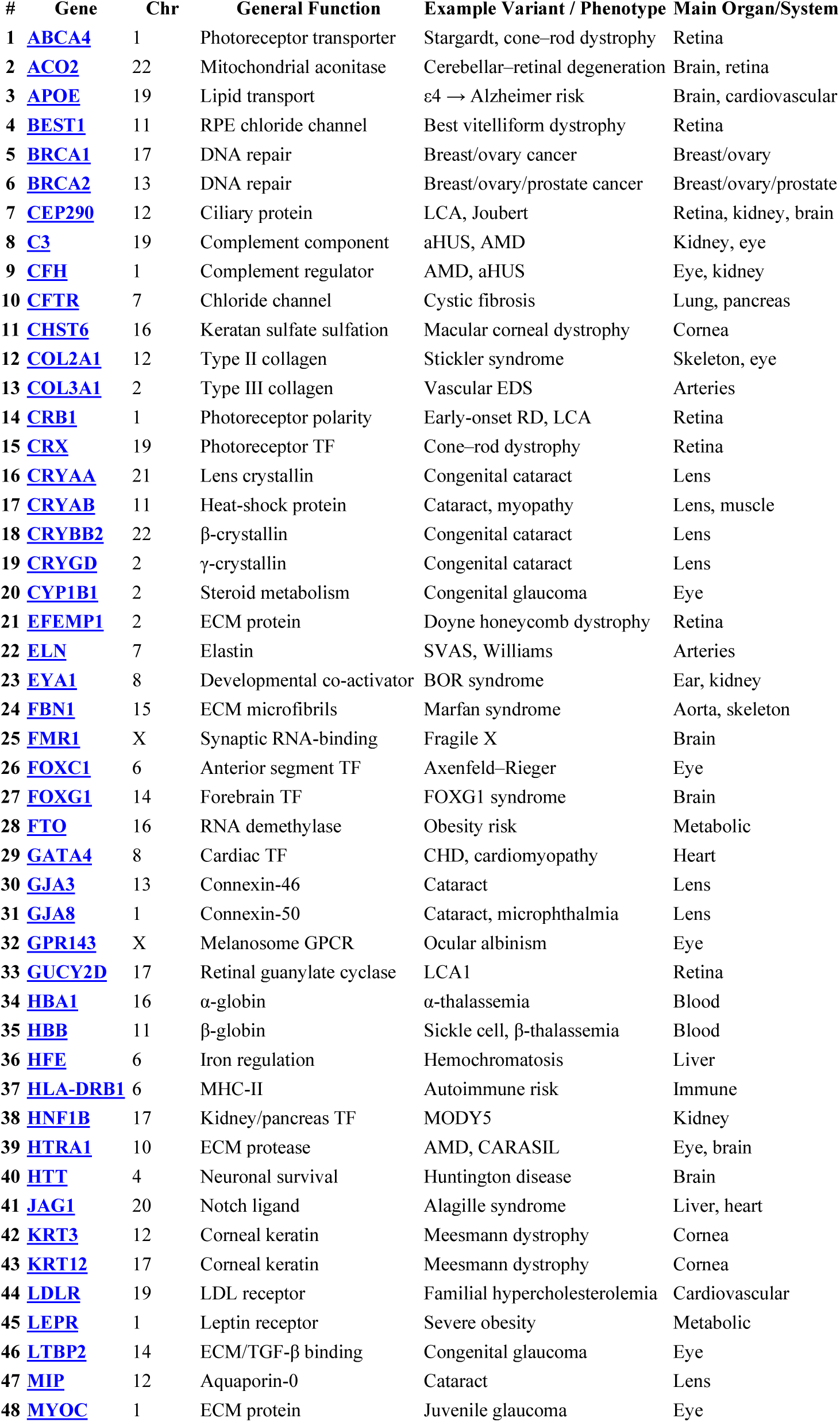
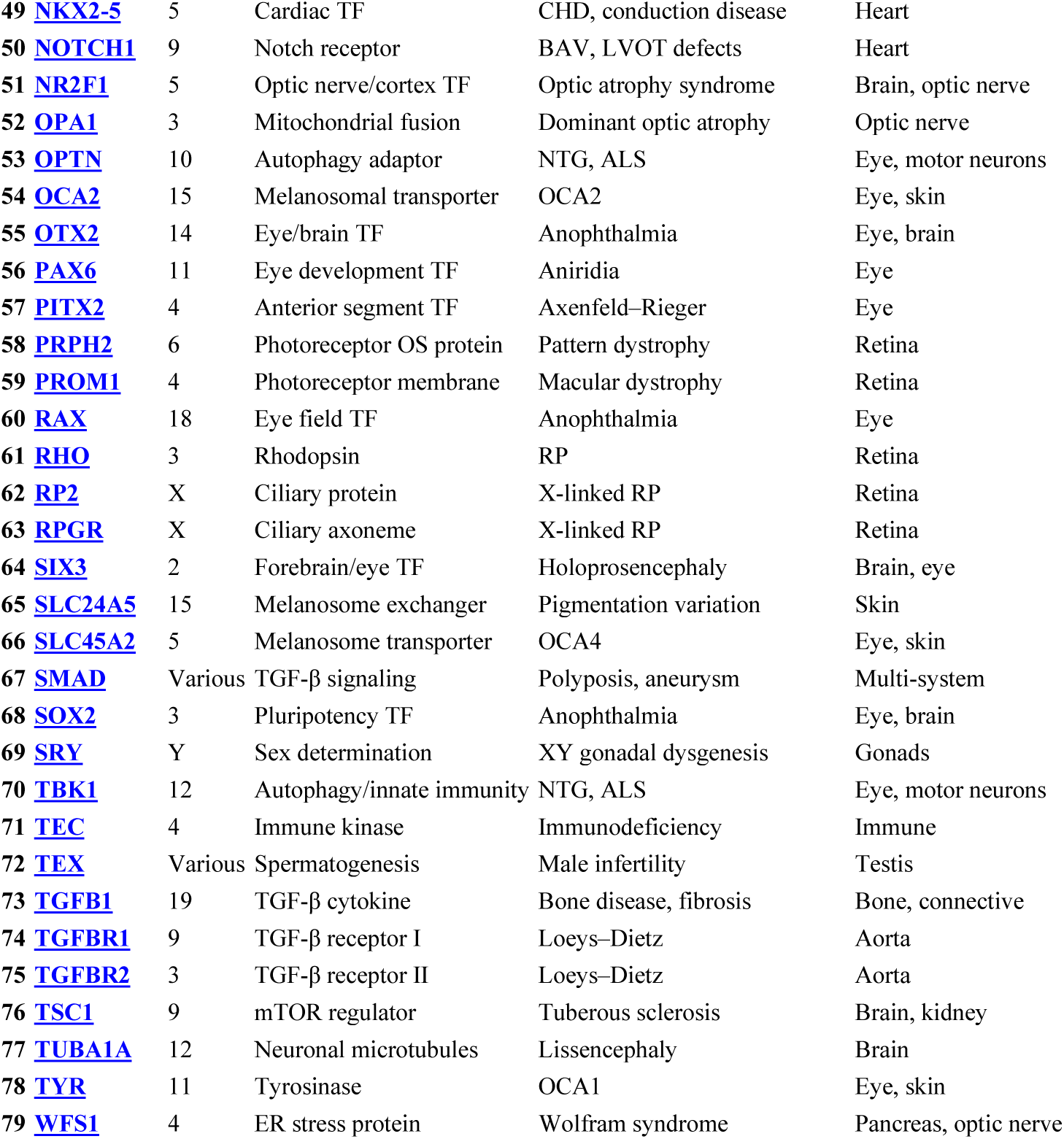
— Nuclear Genes.

**Table B2.**
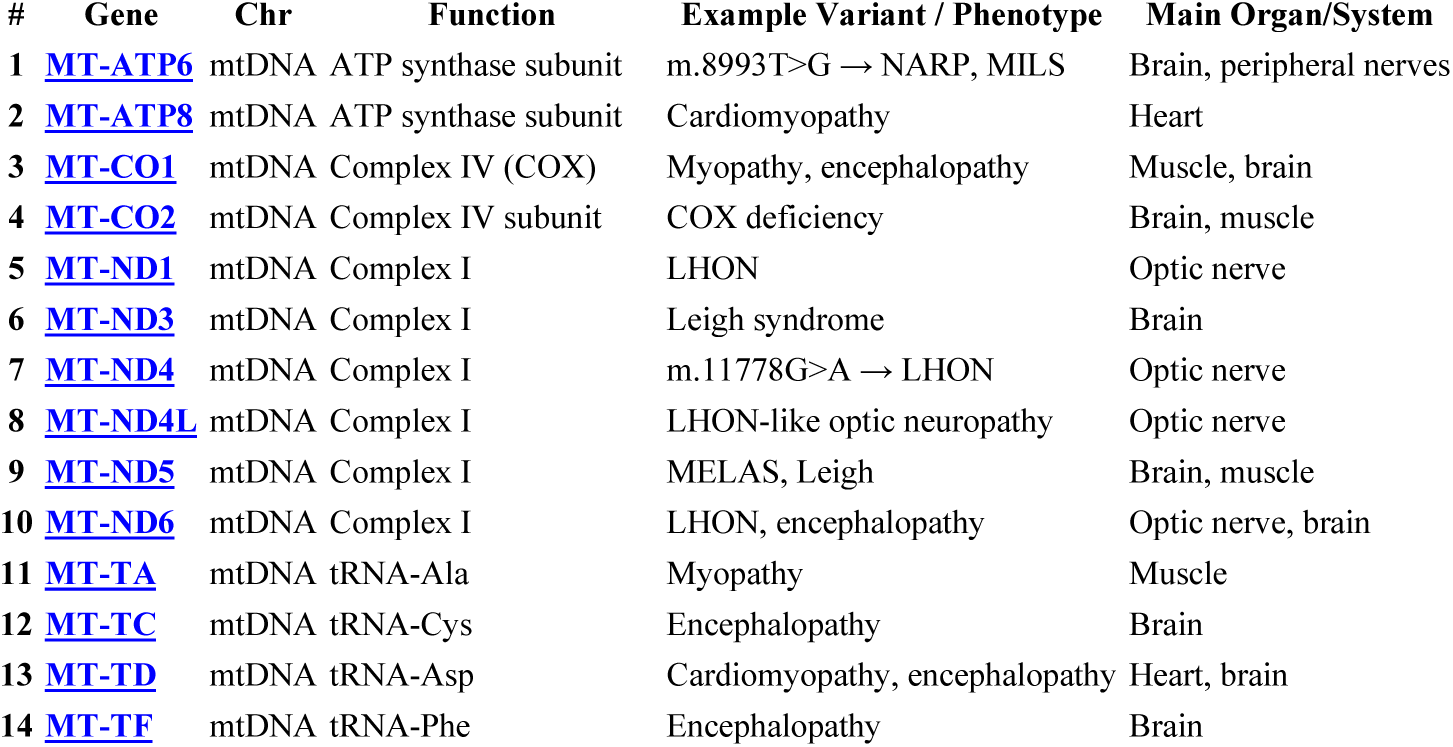
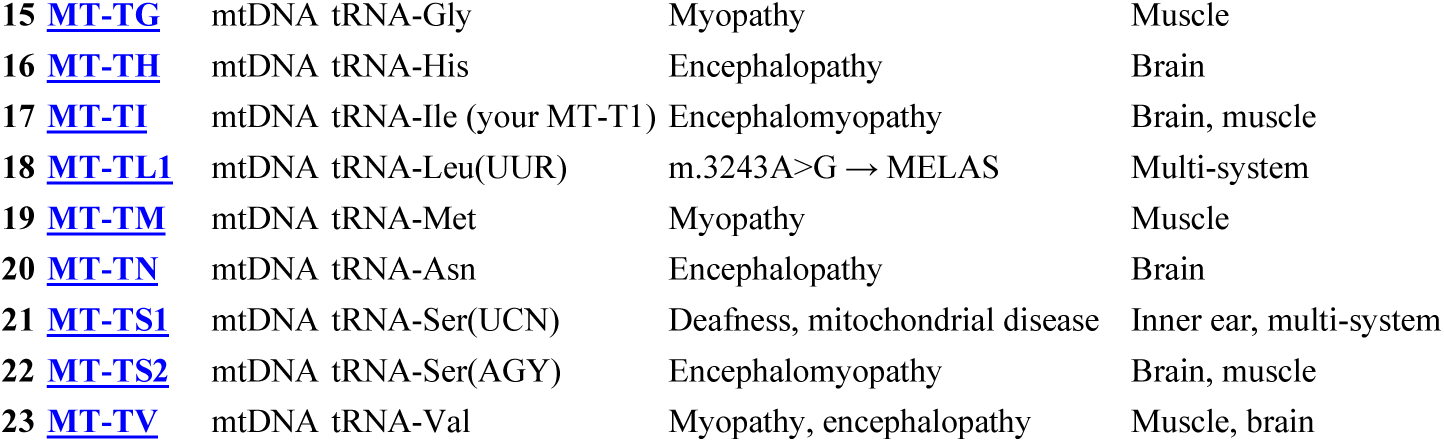
— Mitochondrial Genes.

